# Logomaker: Beautiful sequence logos in python

**DOI:** 10.1101/635029

**Authors:** Ammar Tareen, Justin B. Kinney

## Abstract

Sequence logos are visually compelling ways of illustrating the biological properties of DNA, RNA, and protein sequences, yet it is currently difficult to generate such logos within the Python programming environment. Here we introduce Logomaker, a Python API for creating publication-quality sequence logos. Logomaker can produce both standard and highly customized logos from any matrix-like array of numbers. Logos are rendered as vector graphics that are easy to stylize using standard matplotlib functions. Methods for creating logos from multiple-sequence alignments are also included.

**Availability and Implementation:** Logomaker can be installed using the pip package manager and is compatible with both Python 2.7 and Python 3.6. Source code is available at http://github.com/jbkinney/logomaker.

**Supplemental Information:** Documentation is provided at http://logomaker.readthedocs.io.

## Introduction

Sequence logos provide evocative graphical representations of the functional properties of DNA, RNA, and protein sequences. Logos consist of characters stacked upon one another at a series of integer-valued positions, with the height of each character conveying some type of information about its biological importance. This graphical representation was introduced by Schneider and Stephens (1990) for illustrating statistical properties of multiple-sequence alignments. Although the specific representation they advocated is still widely used, sequence logos have since evolved into a general data visualization strategy that can be used to illustrate many different kinds of biological information (Kinney and McCandlish, 2019). For example, logos can be used to illustrate base-pair-specific contributions to protein-DNA binding energy (Foat *et al.*, 2006), the effects of mutations in massively parallel selection experiments (Liachko *et al.*, 2013), and the saliency footprints of deep neural networks (Shrikumar *et al.*, 2017; Jaganathan *et al.*, 2019).

Making customized logos, however, remains surprisingly difficult. A substantial number of software tools for generating sequence logos have been described, but almost all of these tools limit the kinds of logos one can make and the ways in which those logos can be styled. An important exception is ggseqlogo (Wagih, 2017), which enables the creation of sequence logos within the R programming environment from arbitrary user-provided data. Importantly, ggseqlogo renders logos using native vector graphics, which facilitates post-hoc styling and the incorporation of logos into publication-ready figures. However, similar software is not yet available for Python. Because many biological data analysis pipelines are written in Python, there is a clear need for such logo-generating capabilities.

## Implementation

Logomaker is a flexible Python API for creating publication-quality sequence logos. Logomaker takes as input a matrix (specifically, a pandas data frame) in which columns represent characters, rows represent positions, and values represent character heights (Figure 1A). This enables the creation of logos representing any type of data. The resulting logo is drawn using vector graphics embedded within a standard matplotlib Axes object, thus facilitating a high level of customization as well as incorporation into complex figures that contain other graphical elements. Logomaker provides a variety of options for styling the characters within a logo, including the choice of font, color scheme, vertical and horizontal padding, etc.. Logomaker also enables the highlighting of specific sequences within a logo (Figure 1E), as well as the use of value-specific transparency in logos that illustrate probabilities (Figure 1C). If desired, users can further customize individual characters within any rendered logo.

**Figure 1:**
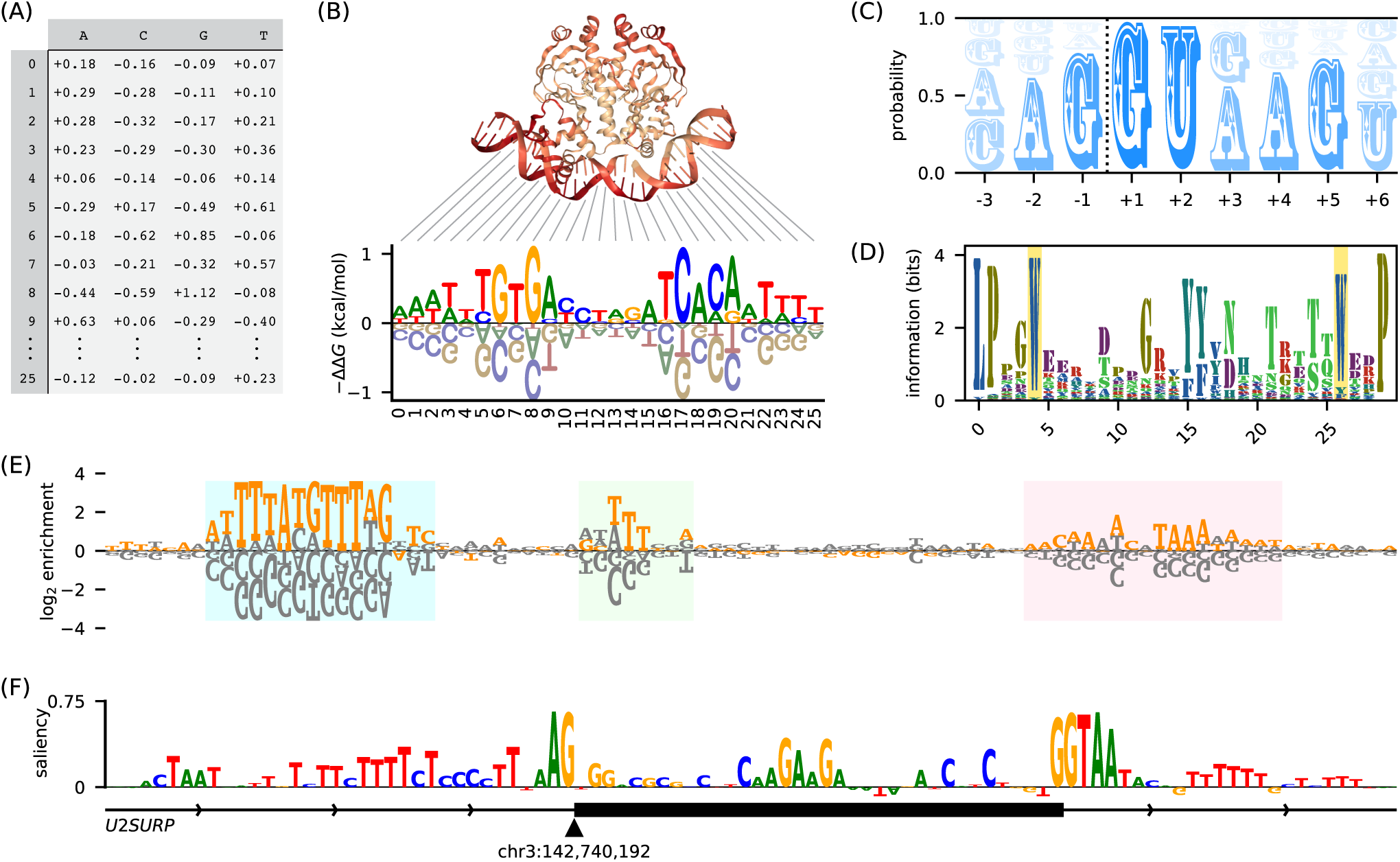
Logomaker logos can represent diverse types of data. (A) Example input to Logomaker. Shown is an energy matrix for the transcription factor CRP; elements of this data frame represent −ΔΔ*G* values contributed by each possible base (columns) at each nucleotide position (rows). Data are from Kinney *et al.* (2010). (B) An energy logo for CRP created by passing the data frame in panel A to Logomaker. The structural context of each nucleotide position is indicated [PDB 1CGP Parkinson *et al.* (1996)]. (C) A probability logo computed from all annotated 5′splices sites in the human genome (Frankish *et al.*, 2019). The dashed line indicates the intron/exon boundary. (D) An information logo computed from a multiple alignment of WW domain sequences [PFAM RP15 (Finn *et al.*, 2014)], with the eponymous positions of this domain highlighted. (E) An enrichment logo representing the effects of mutations within the ARS1 replication origin of *S. cerevisiae*. Orange characters indicate the ARS1 wild-type sequence; highlighted regions correspond (from left to right) to the A, B1, and B2 elements of this sequence (Rao and Stillman, 1995). Data (unpublished; collected by JBK) are from a mutARS-seq experiment analogous to the one reported by Liachko *et al.* (2013). (F) A saliency logo (Shrikumar *et al.*, 2017) representing the importance of nucleotides in the vicinity of *U2SUR* exon 9, as predicted by a deep neural network model of splice site selection. Logo adapted (with permission) from Figure 1D of Jaganathan *et al.* (2019).

Because sequence logos are still commonly used to represent the statistics of multiple sequence alignments, Logomaker provides methods for processing such alignments into matrices that can then be used to generate logos. Multiple types of matrices can be generated in this way, including matrices that represent probabilities (Figure 1C), log odds ratios (Figure 1E), or the information values described by Schneider and Stephens (1990) (Figure 1D). Methods for transforming between these types of matrices are also provided. Finally, Logomaker also supports the creation of matrices (and thus logos) that represent neural network saliency maps (Shrikumar *et al.*, 2017), as in Figure 1F.

## Conclusion

Logomaker thus fills a major need in the Python community for flexible logo-generating software. Indeed, preliminary versions of Logomaker have already been used to generate logos for multiple publications (Belliveau *et al.*, 2018; Wong *et al.*, 2018; Forcier *et al.*, 2018; Nguyen *et al.*, 2018; Barnes *et al.*, 2019; Kinney and McCandlish, 2019). Logomaker is thoroughly tested, has minimal dependencies, and can be installed from PyPI by executing `pip install logomaker` at the command line. A step-by-step tutorial on how to use Logomaker, as well as comprehensive documentation, are available at http://logomaker.readthedocs.io.

## Acknowledgements

We thank Polly Fordyce, William Ireland, David McCandlish, and Bruce Stillman for helpful discussions. We also thank Kyle Farh and Kishore Jaganathan for providing data for the saliency logo in Figure 1F.

